# Chondroitin Sulfate Proteoglycan 4,6 sulfation regulates sympathetic nerve regeneration after myocardial infarction

**DOI:** 10.1101/2022.03.10.483746

**Authors:** MR Blake, DC Parrish, MA Staffenson, S Sueda, W Woodward, BA Habecker

**Author notes:** ^*^**Address for Correspondence**: Beth A. Habecker PhD, Department of Chemical Physiology and Biochemistry, L334, Oregon Health and Science University, 3181 SW Sam Jackson Park Rd, Portland, OR, USA, 97239.

## Abstract

Sympathetic denervation of the heart following myocardial infarction (MI) is sustained by chondroitin sulfate proteoglycans (CSPGs) in the cardiac scar, and denervation predicts risk of sudden cardiac death. Blocking CSPG signaling restores sympathetic axon outgrowth into the cardiac scar, decreasing arrhythmia susceptibility. Axon growth inhibition by CSPGs is thought to depend on the sulfation status of the glycosaminoglycans (CS-GAGs) attached to the core protein. Tandem sulfation of CS-GAGs at the 4^th^ (4S) and 6^th^ (6S) positions of n-acetyl-galactosamine inhibits outgrowth in several types of neurons within the central nervous system, but it is not known if sulfation is similarly critical during peripheral nerve regeneration. We asked if CSPG sulfation prevented sympathetic axon outgrowth. Sympathetic neurite outgrowth across purified CSPGs is restored *in vitro* by reducing 4S with the 4-sulfatase enzyme Arylsulfatase-B (ARSB). Additionally, we co-cultured cardiac scar tissue with sympathetic ganglia *ex vivo* and found that reducing 4S with ARSB restored axon outgrowth to control levels. We examined levels of the enzymes responsible for adding and removing sulfation to CS-GAGs by western blot to determine if they were altered in the left ventricle after MI. We found that CHST15 (4S dependent 6-sulfotransferase) was upregulated, and ARSB was downregulated after MI. Increased CHST15 combined with decreased ARSB suggests a mechanism for production and maintenance of sulfated CSPGs in the cardiac scar. We altered tandem sulfated 4S,6S CS-GAGs *in vivo* by transient siRNA knockdown of *Chst15* and found that reducing 4S,6S restored Tyrosine Hydroxylase (TH) positive sympathetic nerve fibers in the cardiac scar after MI and reduced arrhythmias. Overall, our results suggest that modulating CSPG-sulfation after MI may be a therapeutic target to promote sympathetic nerve regeneration in the cardiac scar and reduce post-MI cardiac arrhythmias.

## Introduction

Sympathetic denervation following myocardial infarction (MI) is well documented in both animal models and in humans, and highly predictive of ventricular arrhythmias and sudden cardiac death(Boogers et al., 2010; Fallavollita et al., 2014; Nishisato et al., 2010). Following an early period of axon degeneration after MI, sympathetic nerves grow back through undamaged myocardium but do not enter the scar due to the presence of chondroitin sulfate proteoglycans (CSPGs) (Gardner & Habecker, 2013). This heterogeneity in sympathetic innervation increases risk for arrhythmias and sudden cardiac death, and blocking sympathetic transmission with beta receptor antagonists or surgical sympathectomy prevents arrhythmias and prolongs life(Herring, Kalla, & Paterson, 2019). However, beta blockers and surgical sympathectomy have unwanted side effects and a therapeutic intervention restoring sympathetic innervation in the cardiac scar by modulating CSPGs may improve health outcomes for patients suffering post-MI arrhythmias.

CSPGs are a diverse family of extracellular matrix proteins modified by chondroitin sulfate (CS) side chains that inhibit nerve regeneration in numerous injury paradigms including MI, spinal cord injury, and traumatic brain injury (Brown et al., 2012; Gardner & Habecker, 2013; Lang et al., 2015; McKeon, Jurynec, & Buck, 1999; Miller & Hsieh-Wilson, 2015; Yi et al., 2012). CSPGs are heterogeneous but all are composed of a core protein that is covalently linked to repeating disaccharide side chains known as glycosaminoglycans (GAGs). While some evidence suggests that the CSPG core protein itself plays a role in regulating axon outgrowth(Dou & Levine, 1994; Ughrin, Chen, & Levine, 2003), the primary effect is thought to occur via the post-translational addition of sulfate to GAGs (Mencio, Hussein, Yu, & Geller, 2021; Miller & Hsieh-Wilson, 2015). CS-GAGs in scar tissues bind CSPG receptors like PTPσ (protein tyrosine phosphatase receptor sigma) on regenerating axons (Coles et al., 2011; Shen et al., 2009). CSPG sulfation is attached to specific locations of the CS-GAG by sulfotransferase enzymes, yielding unique structures that differentially affect axon outgrowth. Specifically, 4-sulfation (4S) and 6-sulfation (6S) of n-acetyl-galactosamine are critical regulators of axon outgrowth (Brown et al., 2012; Gilbert et al., 2005; H. Wang et al., 2008). 4S CS-GAGs are produced by the chondroitin-4-sulfotransferase, CHST11, while 4,6 tandem sulfated GAGs (4S,6S) are produced by a 4S dependent chondroitin-6-sulfotransferase, CHST15 (Miller & Hsieh-Wilson, 2015). 4S CS-GAGs can also be removed by an endogenously expressed 4-sulfatase, Arylsulfatase-B (ARSB) (Figure 1A) (Pearson, Mencio, Barber, Martin, & Geller, 2018; H. Wang et al., 2008). Evidence from spinal cord injury (SCI) and traumatic brain injury (TBI) indicate that tandem sulfated 4S,6S CS-GAGs potently suppress axon outgrowth(Brown et al., 2012; Gilbert et al., 2005). Removal of all CS-GAGs from cardiac scar tissue by treatment with the enzyme chondroitinase ABC (chABC) restores sympathetic axon outgrowth *in vitro* (Gardner & Habecker, 2013) but it remains unknown whether CSPG sulfation is critical to inhibit regeneration of peripheral nerves.

**Figure 1:**
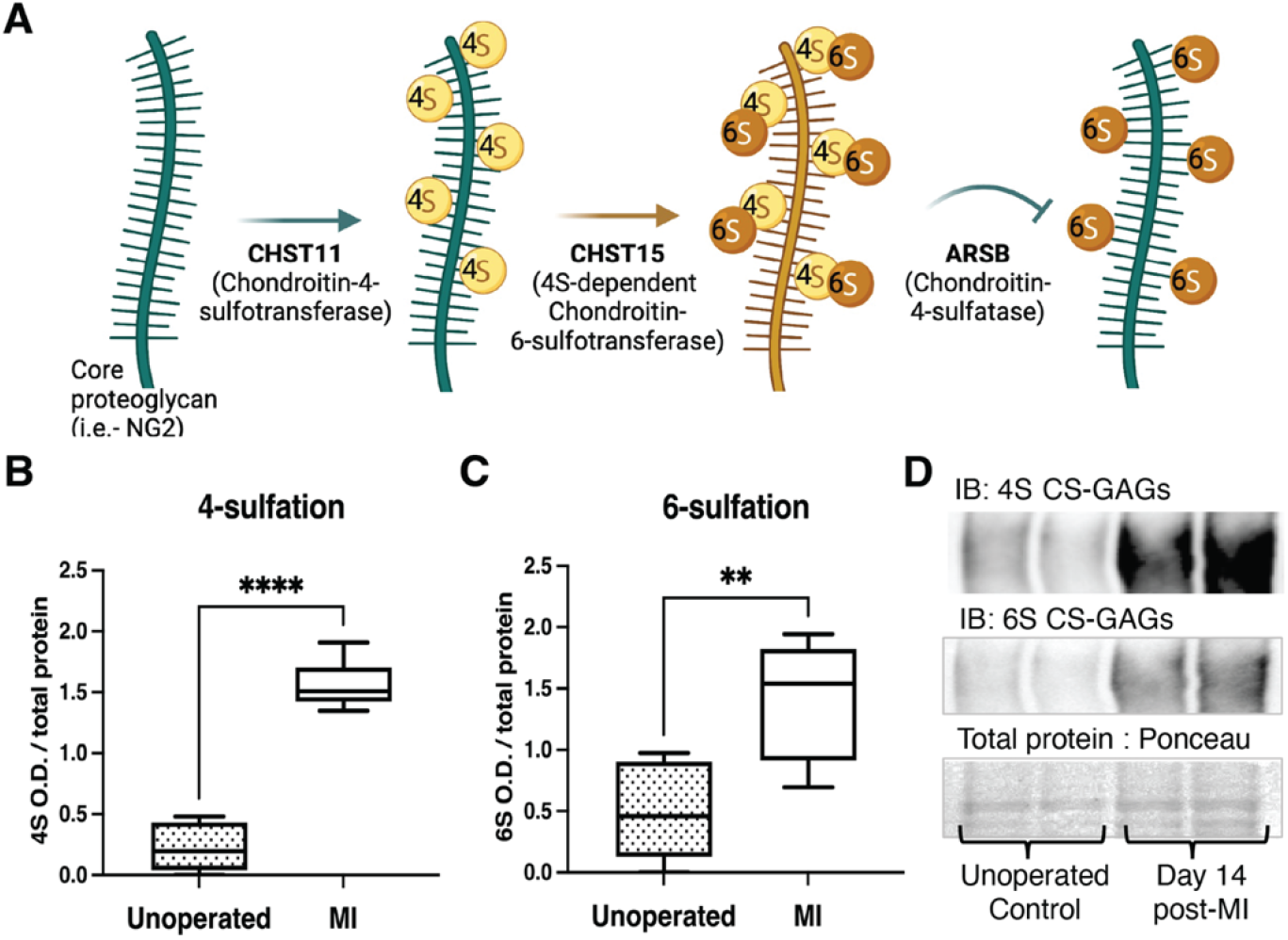
CSPG 4,6 Sulfation increases in the cardiac scar after MI. **(A)** CSPG-sulfation patterning schematic with key enzymes. **(B)** 4-sulfation (4S CS-GAGs) or **(C)** 6-sulfation (6S CS-GAGs) quantification assessed by western blot in the healthy myocardium (unoperated) or cardiac scar 14 days after MI (MI). Quantification of n=6 animals per treatment group, statistics; student t-test (Welch’s test), **p-value<0.01, ****p-value<0.0001 **(D)** Example blot images for 4S CS-GAGs, 6S CS-GAGs, and total protein from two unoperated and two MI animals.

Here we show that 4,6 tandem sulfated CSPGs are enriched in the cardiac scar after MI, and that this sulfation is important for preventing nerve regeneration in the heart. Sulfation-related enzymes are altered in the heart after MI, and reducing sulfation of CS-GAGs by transient siRNA knockdown of *Chst15* promotes reinnervation of the cardiac scar. Reinnervation decreases isoproterenol-induced arrhythmias.

## Results

### 4,6-sulfation of CS-GAGs increased in the heart after MI

To establish whether CSPG sulfation occurs after MI we used antibodies specific to 4S and 6S CS-GAGs (Yi et al., 2012) to compare CSPG sulfation in unoperated left ventricle to day 14 post-MI scar tissue. Day 14 represents a time point when scar tissue is relatively stable and sympathetic nerves are excluded from entering the scar. Myocardial infarction led to increased 4,6 sulfation of CSPGs in the cardiac scar (Figure 1B-D).

### Reducing 4-sulfation with ARSB promotes sympathetic neurite outgrowth in vitro

To test whether 4-sulfation of CS-GAGs prevented neurite outgrowth across CSPGs we enzymatically removed 4S from purified CSPGs using the 4-sulfatase, ARSB. An effective ARSB dose was identified in pilot studies to remove 4S while leaving 6S intact (Figure S1). Sympathetic neurons were then grown on CSPGs treated with vehicle or ARSB. Sympathetic neurite extension across untreated CSPGs was suppressed 40 hours post-plating compared to laminin alone, while ARSB treatment restored neurite outgrowth in a dose-dependent manner (Figure 2A,B). These results indicate that 4-sulfation of CS-GAGs inhibits sympathetic neurite extension.

**Figure 2:**
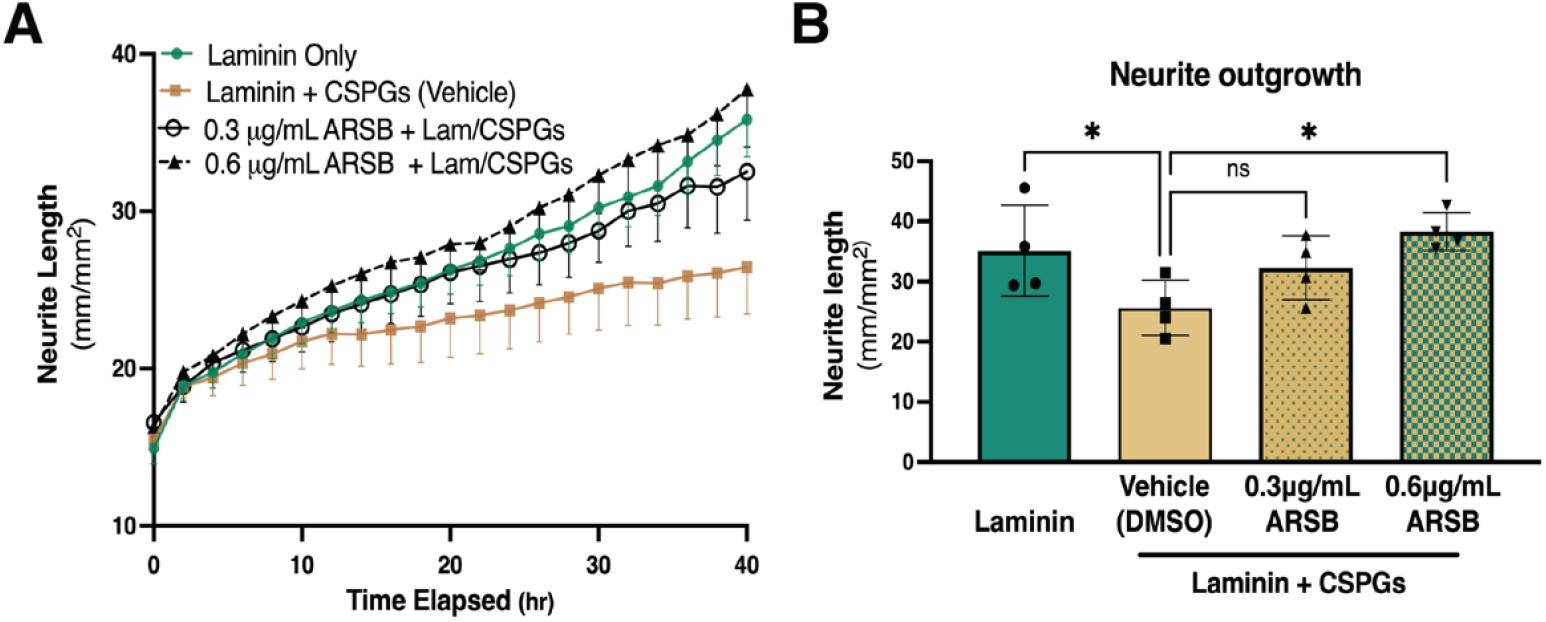
Reducing CS-GAG 4-sulfation promotes sympathetic neurite outgrowth *in vitro*. **(A)** Example of neurite outgrowth experiment with ARSB. Data are mean neurite-length ± SD at 9 locations per well and 3 wells per condition. **(B)** Removal of 4-sulfation with ARSB restores neurite outgrowth to control levels. Quantification of dissociated sympathetic neurite length at 40 hours post-plating on indicated plate coatings. Statistics; one-way ANOVA, (Dunnett’s post-test). All comparisons made to neurite outgrowth on Laminin + CSPG coated wells, ns - not significant, *p<0.05, n=4 experiments.

### Reducing 4-sulfation of cardiac scar tissue with ARSB restores sympathetic axon outgrowth ex vivo

We treated cardiac scar tissue with ARSB to test whether removing 4-sulfation from CS-GAGs in cardiac scar tissue could restore sympathetic axon outgrowth in explant co-cultures. Ganglion explants co-cultured (Figure 3A) with unoperated left ventricle (Figure 3B) displayed uniform axon outgrowth in all directions while explants cultured alongside cardiac scar tissue (10-14 days post-MI) exhibited shorter axons growing in the direction of the scar tissue (Figure 3C). Interestingly, ARSB treatment (0.6μg/mL) of scar tissue fully restored axon outgrowth towards the scar (Figure 3D), while growth away from the scar tissue was normal in all conditions (Figure 3E). These data suggest that reducing 4-sulfation of CS-GAGs in the cardiac scar is sufficient to enable sympathetic axon regeneration.

**Figure 3:**
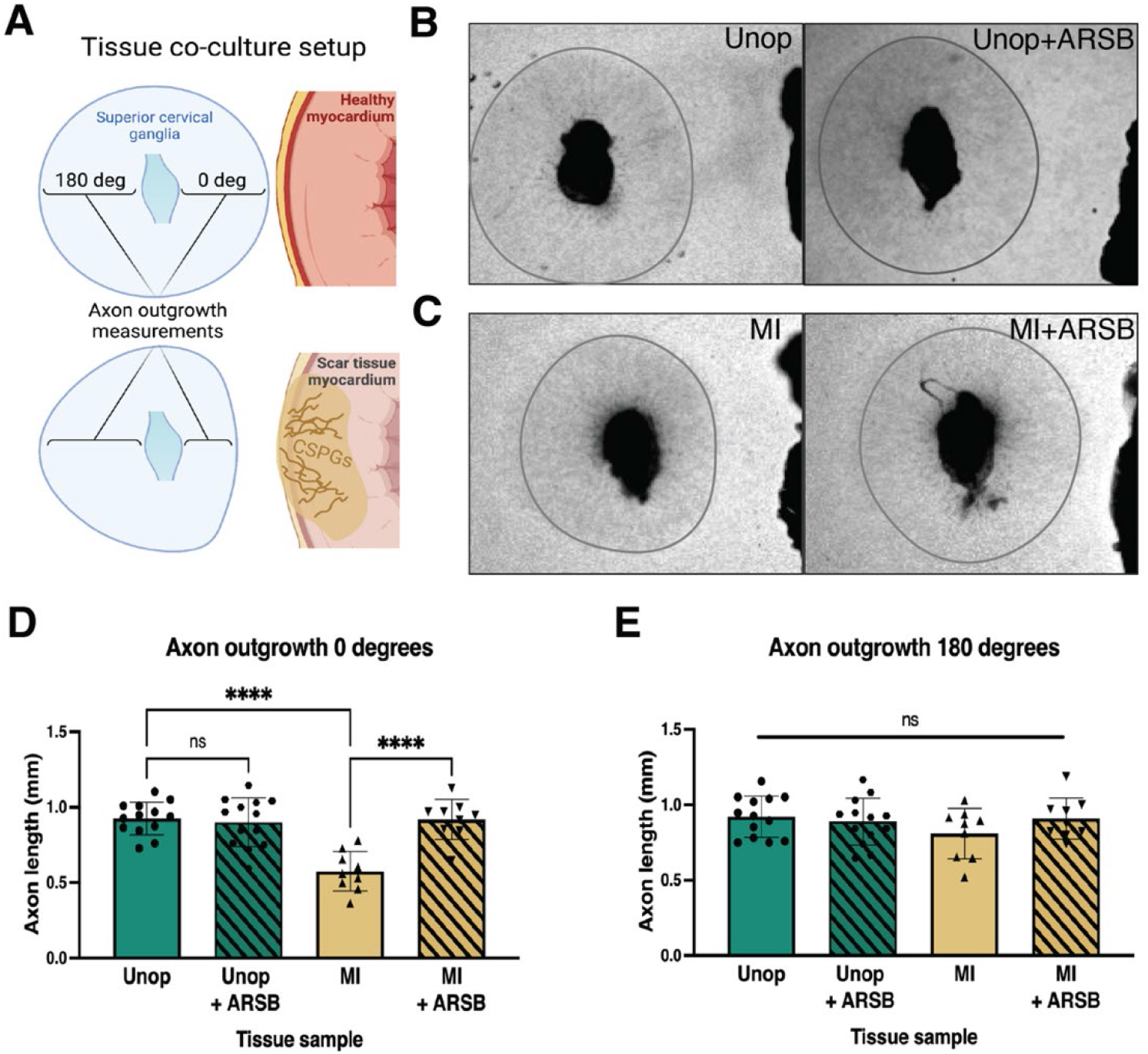
Reducing CS-GAG 4-sulfation in cardiac scar tissue *ex vivo* restores sympathetic axon outgrowth. **(A)** Explant co-culture schematic. Ganglion axon extension towards (0 degree) and away from (180 degree) myocardium was measured. **(B)** Example images of ganglia axon outgrowth in the presence of healthy LV tissue (Unop) or **(C)** cardiac scar tissue (MI) treated with or without ARSB. Gray circle approximates boundary of axon extension 48 hours after plating. **(D)** Quantification of ganglion axon outgrowth towards myocardium of either healthy tissue (Unop) or scar tissue (MI) treated with vehicle (5% DMSO) or ARSB (0.6μg/mL). **(E)** Quantification of ganglion axon outgrowth away from myocardium of either healthy tissue (Unop) or scar tissue (MI) treated with vehicle (5% DMSO) or ARSB (0.6μg/mL). Statistics for D, E; one-way ANOVA (Dunnett’s post-test), comparisons made to vehicle treated MI tissue; ns - not significant, p<0.0001, n=13 animals control tissue and n=9 MI tissue.

### Expression of CSPG sulfation enzymes is altered after MI

To understand the mechanisms by which CS-GAG sulfation increases after MI we examined protein levels of three critical CSPG sulfation enzymes, comparing post-MI expression to unoperated left ventricle. We examined chondroitin-4-sulfotransferase CHST11, 4S-dependent chondroitin-6-sulfotransferase CHST15 (4,6 tandem sulfation enzyme), and 4-sulfatase ARSB (Figure 1A). CHST11 levels decreased significantly within 24 hours, persisting until day 7 (Figure 4A,D), while CHST15 was increased significantly at days 7 and 14 post-MI (Figure 4B,D). ARSB levels decreased significantly by day 3 and remained low through day 14 (Figure 4C,D).

**Figure 4:**
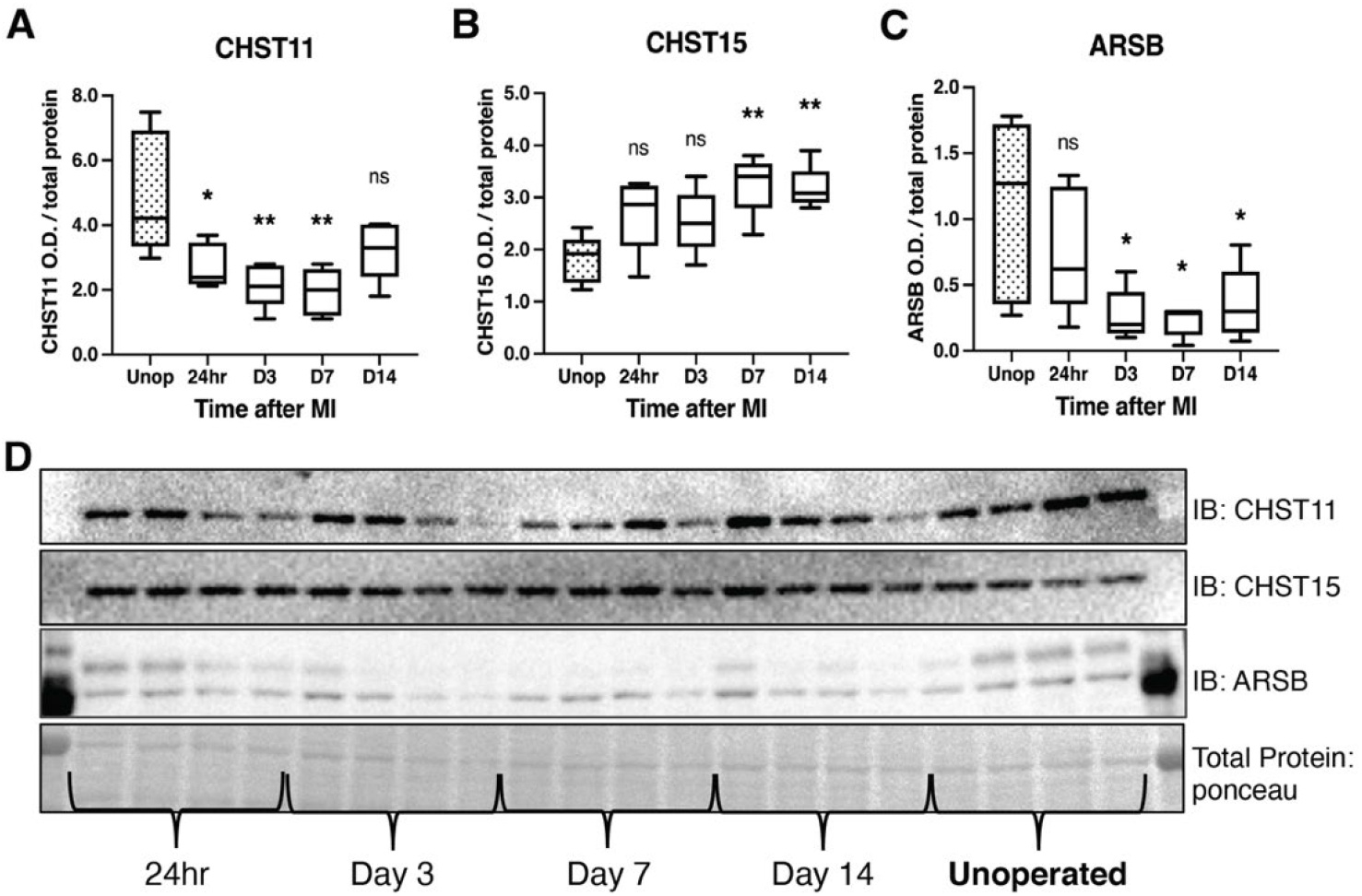
CSPG-sulfation enzyme expression is altered after MI. Western blot quantification of protein expression for **(A)** chondroitin-4-sulfotransferase (CHST11), **(B)** chondroitin-6-sulfotransferase (CHST15) and, (**C)** 4-sulfatase (ARSB) protein expression in control left-ventricle (Unop), or in the days after MI (24hr, D3, D7, or D14). Statistics; one-way ANOVA (Dunnett’s post-test); ns-not significant, *p-value<0.05, **p-value<0.01, comparisons to unoperated tissue, n=5 animals per group. **(D)** Example western blot images for CSPG-sulfation enzymes.

In light of protein changes in CSPG sulfation enzymes after MI, we examined CS-GAG sulfation at these same time points. 4S and 6S CS-GAGs are significantly increased in the infarct by day 7 post-MI extending through day 14 (Figure 5A,B,E). We also asked if the CSPG core protein NG2 (also called CSPG4) was altered after MI, and found that NG2 was increased in the infarct by 7 days after MI (Figure 5C,E). Thus, core proteins and sulfation of CS-GAG side chains are both in greater abundance. We chose to look at NG2 because it was prominently expressed in a recent glycoproteomics analysis of cardiac scar tissue whereas another commonly studied core proteoglycan, Aggrecan, was not detected (Tian et al., 2014). In an effort to better understand how this process may be coordinated, we examined the expression of a key wound healing regulator, Galectin-3, which has also been connected to CSPG sulfation and ARSB expression (Bhattacharyya et al., 2020; Bhattacharyya, Feferman, Terai, Dudek, & Tobacman, 2017; Bhattacharyya, Feferman, & Tobacman, 2014; Bhattacharyya et al., 2015). Galectin-3 was increased significantly by 24 hours after MI, which may contribute to altered expression of CSPG sulfation enzymes at later time points (Figure 5D,E). These data suggest that increased CHST15 and decreased ARSB expression contribute to increased 4,6 sulfation of CS-GAGs observed by day 7 post-MI.

**Figure 5:**
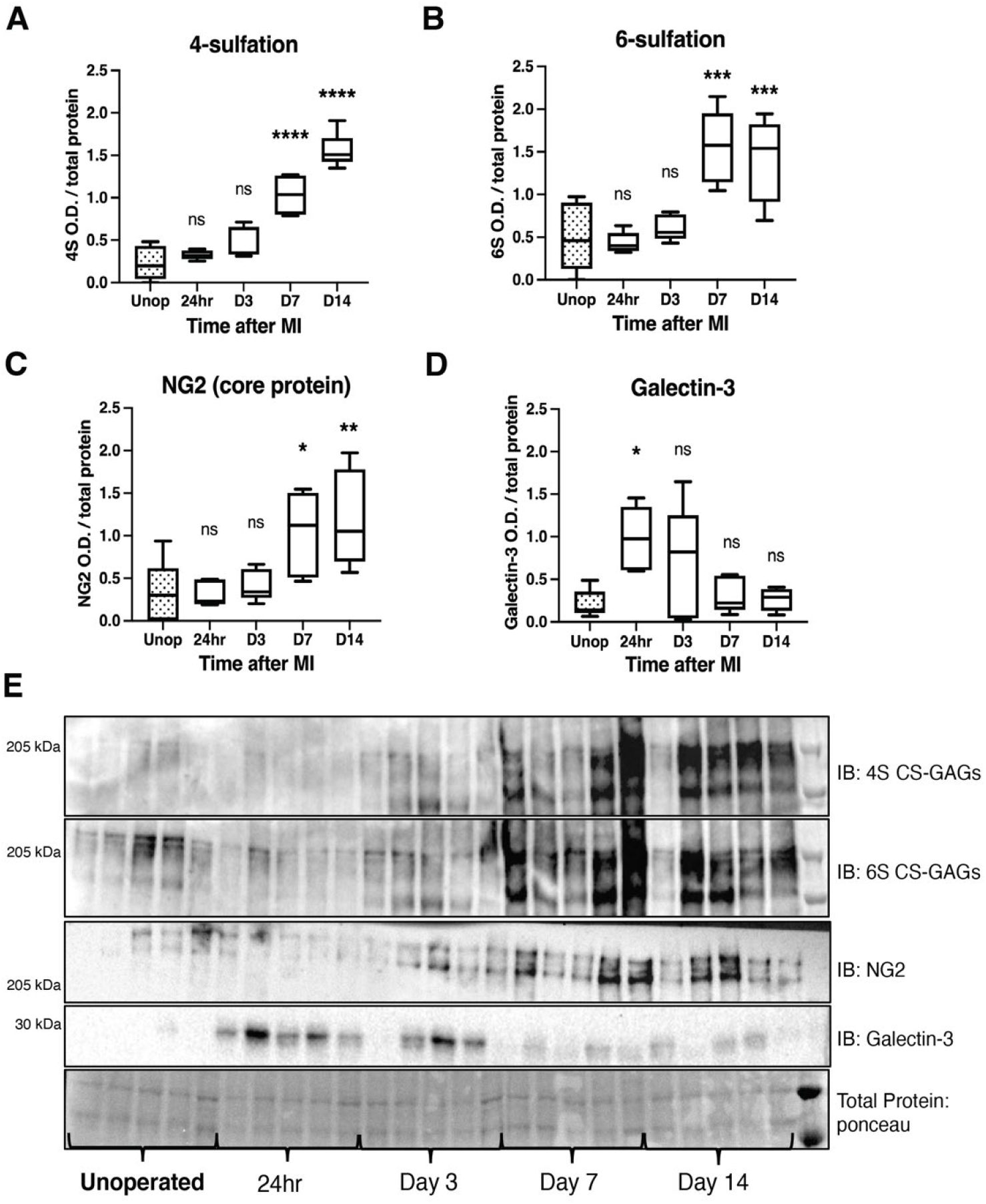
Time course of CSPG sulfation and core protein expression after MI in cardiac scar tissue. Western blot quantification of **(A)** 4-sulfation (4S CS-GAGs), **(B)** 6-sulfation ( 6S CS-GAGs), **(C)** NG2 core protein, and **(D)** Galectin-3 in the days after MI. Statistics; one-way ANOVA (Dunnett’s post-test), ns-not significant, *p-value<0.05, **p-value<0.01, ***p-value<0.001, ****p-value<0.0001, comparisons to unoperated tissue, n=5 animals per group. **(E)** Example western blot images of A-D.

### siRNA knockdown of Chst15 reduces 4,6 tandem sulfation of CSPGs and restores sympathetic axons in the cardiac scar after MI

To determine if CSPG sulfation suppresses sympathetic nerve regeneration after MI we decreased 4,6-tandem sulfated CS-GAGs *in vivo* by reducing the expression of the *Chst15* gene using silencing RNA (siRNA). After identifying siRNA that decreased *Chst15* mRNA in myoblast-like C2C12 cells (Figure S2A, S2B), we tested the efficacy of our *Chst15* siRNA *in vivo*. Intravenous delivery of 100μg si*Chst15* reduced CHST15 protein compared to non-targeting control siRNA (Figure S2C). Mice were then treated with 100μg si*Chst15* on days 3, 5, and 7 post-MI, leading to a significant reduction in 4S,6S CS-GAGs. Specifically, si*Chst15* reduced the amount of 6S present 10 days post-MI compared to non-targeting siRNA control (Figure 6A). By 10 days after MI, CHST15 protein levels had returned to normal suggesting that transient knockdown of CHST15 is sufficient to alter 4,6 sulfation of CS-GAGs (Figure S2D,E). Interestingly, si*Chst15* knockdown led to increased NG2 core protein expression compared to non-targeting siRNA control (Figure 6C). The sympathetic neuron marker Tyrosine Hydroxylase (TH), was increased significantly in cardiac scar tissue from si*Chst15* treated animals compared to non-targeting siRNA control animals (Figure 6D), suggesting successful sympathetic reinnervation of the cardiac scar. Neuronal norepinephrine (NE) content was not increased (Figure S2F), consistent with previous studies showing regulation of NE synthesis and reuptake by inflammatory cytokines (Parrish et al., 2010).

**Figure 6:**
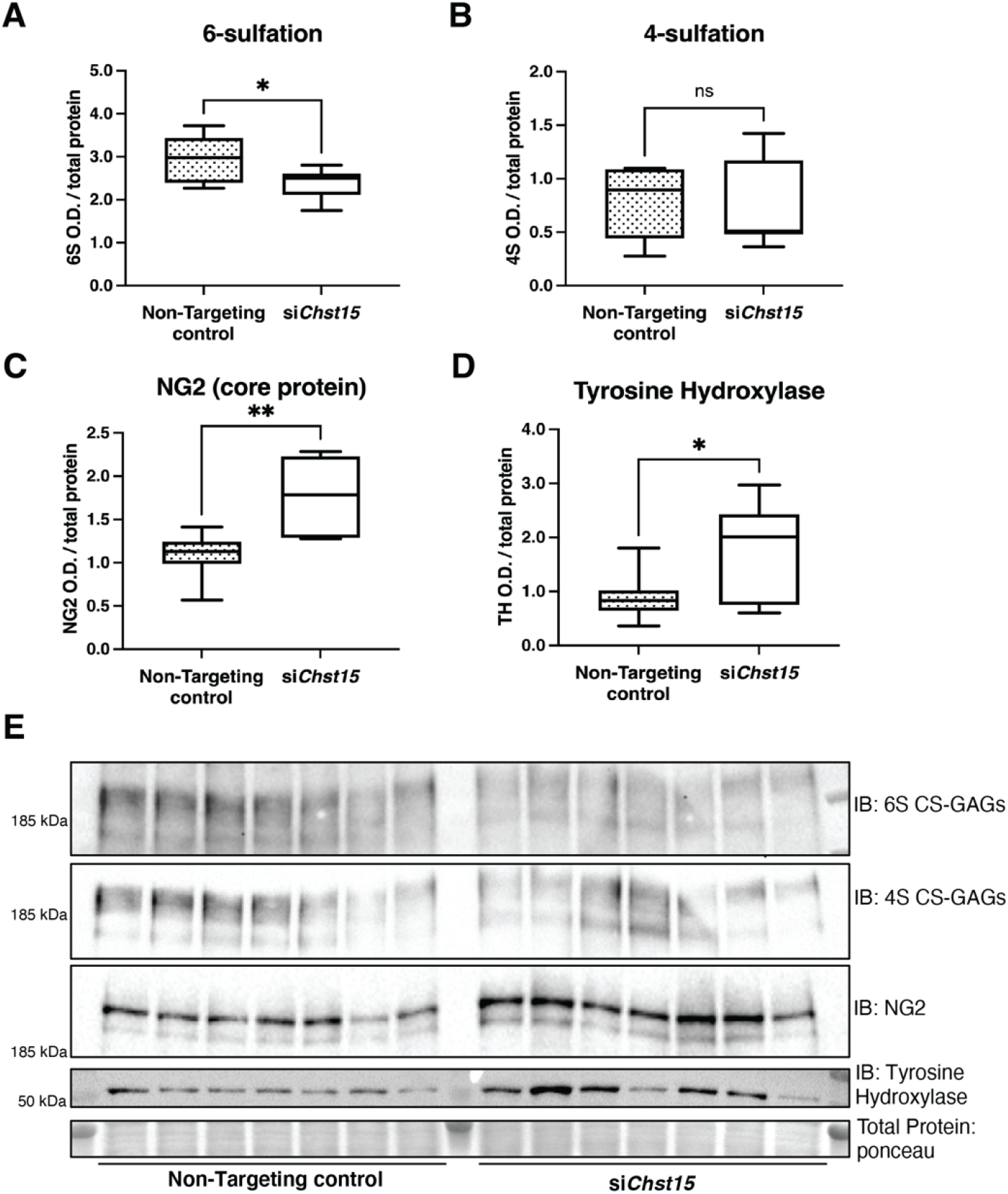
Reducing CS-GAG 4,6 tandem-sulfation *in vivo* promotes sympathetic nerve regeneration into the cardiac scar after MI. Western blot quantification of **(A)** 6-sulfation (6S CS-GAGs), **(B)** 4-sulfation (4S CS-GAGs), **(C)** NG2 core protein and, **(D)** the sympathetic neuron marker Tyrosine Hydroxylase (TH) in the cardiac scar after transient *Chst15* knockdown or treatment with a non-targeting siRNA control. Tissue was collected 10 days after MI, n=7 animals per group; student t-test (Welch’s test), ns - not significant, *p-value<0.05, **p-value<0.01. **(E)** Western blot images of A-D.

To ensure that reinnervation of the cardiac scar occurred after *Chst15* knockdown we examined the cardiac scar (labeled by fibrinogen) for TH positive sympathetic nerve fibers by immunohistochemistry (IHC). IHC analysis in animals treated with a non-targeting siRNA control showed clear denervation of the infarct (Figure 7A) compared to the peri-infarct region adjacent to the scar (Figure 7B). Animals treated with an siRNA targeting *Chst15* had restored TH positive fibers in the infarct (Figure 7C) at the same innervation density as the peri-infarct region (Figure 7D). TH innervation density was significantly higher in the infarct of *Chst15* targeted animals compared to non-targeting siRNA treated animals (Figure 7E). Cardiac scar size was examined between the two groups and no significant difference existed (Figure 7G,H). To examine whether restoration of nerves reduced arrhythmia susceptibility we examined arrhythmias by ECG after administration of the *ββ*-agonist Isoproterenol(Gardner et al., 2015). Restoration of nerves with *Chst15* siRNA treatment reduced arrhythmias when compared to non-targeting siRNA control treated animals (Figure 7F). These results indicate that reducing 4,6 tandem sulfated CS-GAGs in the cardiac scar after MI promotes sympathetic nerve regeneration back into the cardiac scar and reduces arrhythmia susceptibility after MI.

**Figure 7:**
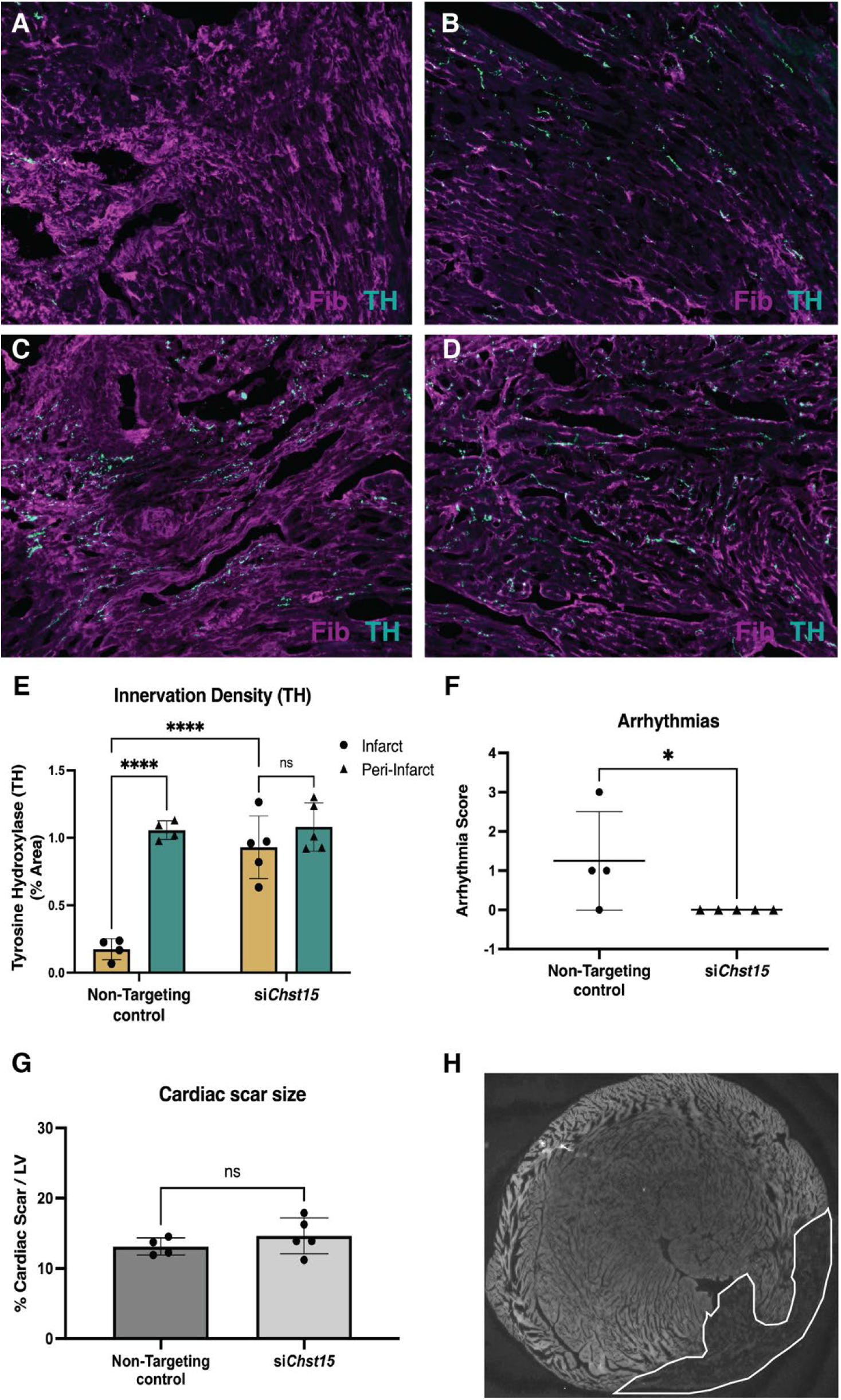
Reducing 4,6 tandem-sulfation of CS-GAGs *in vivo* promotes sympathetic nerve regeneration into the cardiac scar. Fibrinogen was used to label scar (magenta) and TH was used to label sympathetic neurons (cyan). Example image of **(A)** denervated infarct in non-targeting control treated animal versus **(B)** normal density of TH+ fibers in the peri-infarct region adjacent to the scar. Systemic delivery of siRNA against *Chst15* post-MI restored TH+ fiber density in **(C)** the cardiac infarct similarly to **(D)** the peri-infarct region. **(E)** Quantification of TH innervation density (% area, 20x field of view) 10 days after MI. n=4 animals for non-targeting controls and n=5 for *Chst15* siRNA treated animals. Two-way ANOVA, Tukey’s post-test to compare all groups, select comparisons shown, ns - not significant, ****p-value<0.0001. **(F)** Arrhythmia scores based on the most severe arrhythmia observed in each heart after injection of isoproterenol and caffeine (0=no PVCs, 1=single PVCs, 2=bigeminy or salvos, 3= non-sustained ventricular tachycardia) See methods for details. Treatment with siRNA against *Chst15* reduced arrhythmias compared to non-targeting controls. n=4 animals for non-targeting controls and n=5 animals for *Chst15 siRNA*. Statistics; student t-test (Mann-Whitney test), *p<0.05. **(G)** Cardiac scar size assessed as a % area of total left ventricle (LV) area was unaltered in *Chst15* siRNA treated animals compared to non-targeting controls. Quantification of n=4 animals for non-targeting controls and n=5 for *Chst15 siRNA* treated animals. Statistics; student t-test (Welch’s test), ns - not significant. **(H)** Example 2x image of cardiac scar in *Chst15* siRNA treated heart using autofluorescence, absence of autofluorescence indicates region of the infarct, outlined in white.

## Discussion

This study investigated the role of CS-GAG side chain sulfation in preventing sympathetic axon regeneration in the heart after MI. We found that the CSPGs present in the cardiac scar after ischemia-reperfusion were enriched with sulfation at the 4 and 6 positions of CS-GAGs, which is often referred to as chondroitin sulfate E (CS-E) (Miller & Hsieh-Wilson, 2015). CS-E inhibits the outgrowth of several types of neurons in the CNS (Brown et al., 2012; Gilbert et al., 2005), and our data indicate that it also prevents growth of sympathetic axons, since decreasing either 4 or 6 sulfation allowed axon regeneration. Nerve regeneration was confirmed by histology and by expression of the neuronal protein Tyrosine Hydroxylase in cardiac scar tissue. Reinnervation throughout the left ventricle decreased isoproterenol induced arrhythmias, consistent with previous studies targeting the CSPG receptor PTPσ(Gardner et al., 2015).

The core protein NG2 is an important source of CSPGs in the cardiac scar (Tian et al., 2014), and there is evidence that the NG2 core protein can inhibit sensory neurite extension independent of CS sulfation in certain contexts (Dou & Levine, 1994; Ughrin et al., 2003). Modulating the cardiac scar is of interest as a strategy to improve outcomes after MI, but removing matrix proteins presents a challenge since the scar plays an important role in maintaining structural integrity. Given the critical role of CSPG sulfation in other contexts (Brown et al., 2012; Gilbert et al., 2005; Miller & Hsieh-Wilson, 2015; Pearson et al., 2018; H. Wang et al., 2008; Yi et al., 2012) we postulated that this might present an intervention point. While our data indicate that NG2 expression increases following MI, we found that removing or preventing sulfation of CS-GAG chains was sufficient to allow axon regeneration into the cardiac scar. Thus, NG2 prevents sympathetic axon regeneration in the heart exclusively through 4,6 tandem sulfation of CS side chains.

Increased 4,6 sulfation of CS-GAGs coincided with increased expression of the 4S-dependent 6-sulfotransferase CHST15, which is the final enzyme in the production of CS-E, and depletion of the sulfatase ARSB which removes 4 sulfation. These changes in enzyme expression provide a potential mechanism for the shift to highly sulfated GAG side chains after MI. Regulation of these proteins is also linked in the context of traumatic brain injury and other diseases with a downregulation of ARSB leading to increased CHST15 (Bhattacharyya et al., 2020; Bhattacharyya et al., 2017; Bhattacharyya et al., 2014; Bhattacharyya et al., 2015). Hypoxia reduces ARSB activity, increasing the abundance of 4 sulfated CS-GAGs which enables Galectin-3 mediation of transcriptional changes (Bhattacharyya et al., 2014). Therefore, we asked if galectin-3 expression was altered in the heart after ischemia-reperfusion. To our surprise, we found that Galectin-3 increased before changes were observed expression of sulfation-related enzymes. This suggests that increased Galectin-3 may be an early driver of the accumulating highly sulfated CS-GAGs observed by day 7 post-MI.

The inhibitory effects of CS-E on axon outgrowth in certain neuronal subtypes led many to hypothesize that reducing CHST15 activity would restore nerve growth across CSPGs (Brown et al., 2012). To that end at least one group has even generated a novel small molecule to inhibit CHST15 and promote nerve growth (Cheung et al., 2017); However, since no commercially available inhibitor exists to our knowledge, we sought a different method. Targeting *Chst15* gene expression with siRNA decreased 4,6 tandem sulfation of CS-GAGs and allowed nerve regeneration into the cardiac scar. We were inspired to test *chst15* siRNA by a colitis study in mouse showing that siRNA knockdown of *chst15* reduced CS-E (Suzuki et al., 2016; Suzuki et al., 2017). More importantly, *Chst15* siRNA has been tested in a Phase 1 Clinical for Crohn’s Disease patients with active mucosal lesions, and a Phase I/IIa trial for patients with pancreatic cancer(Tsuchiya et al., 2021), and has a good safety profile (Suzuki et al., 2017). Thus, siRNA knockdown of *Chst15* is potentially translatable to large animals and eventually human studies. We treated mice on days 3, 5, and 7 post-MI with siRNA as we wanted to intervene while the cardiac scar was forming. We assessed nerve regeneration into the scar and CSPG sulfation on day 10, which is 3 days after the final injection of siRNA. Three days after the last siRNA injection, CHST15 levels were similar in non-targeting siRNA and *Chst15* siRNA hearts despite a significant reduction in 6-sulfation and reinnervation of the infarct in *Chst15* siRNA treated animals. These data indicate that a transient decrease in CHST15 protein and depletion of CS-GAG sulfation is sufficient to allow nerve restoration in the cardiac scar where NGF is abundant. We also attempted to use an adeno-associated virus 9 (AAV9) virus to elevate ARSB expression after MI, since explant studies showed that ARSB could promote axon outgrowth in the presence of scar-derived CSPGs. However, we were unable to increase ARSB expression in the heart, although the AAV9-GFP (green fluorescent protein) control generated good expression in myocardium. ARSB is an FDA approved therapeutic (Naglazyme™) to treat Mucopolysaccharidosis IV disease (Harmatz et al., 2005; Harmatz et al., 2004; Munoz-Rojas et al., 2010), which may be of use in the therapeutic context of the post-MI heart. Ultimately this work demonstrates the critical nature of 4, 6 sulfation of CS-GAGs in preventing reinnervation of the heart after MI and provides two strategies to promote reinnervation by modulating CSPG sulfation.

## Materials and Methods

### Western blotting

Cardiac scar tissue from the left ventricle was dissected at 24 hours, 3 days, 7 days and 14 days following MI, and control left ventricle tissue from unoperated animals. Heart tissue was pulverized in a glass douncer in NP40 lysis buffer [50mM Tris (pH 8.0), 150mM NaCl, 2mM EDTA, 10mM NaF, 10% glycerol, and 1% NP-40] containing complete protease inhibitor cocktail (Roche), phosphatase inhibitor cocktails 2 and 3 (Sigma). Lysates sat on ice for 30min with intermittent vortexing. Lysates were centrifuged (13k rpm, 10min, 4°C) and resolved on 4-12% Bis-Tris gradient gel (3-8% Tris-Acetate gel for Figure 5 sulfation studies) by SDS/PAGE, transferred to nitrocellulose membrane (GE Life Sciences), blocked in 5% nonfat milk, probed with either CHST11 (1:500; Invitrogen: PA5-68129), CHST15 (1:1000; Proteintech: 14298-1-AP), ARSB (1:500; Proteintech: 13227-1AP), NG2/CSPG4 (1:1000; Millipore: AB5320), Tyrosine Hydroxylase (1:1000; Millipore: AB1542), or Galectin-3 (1:500; Abcam: AB2785) then probed with goat anti-rabbit (or mouse) HRP-conjugated secondary antibody (1:10,000; Thermo), and detected by chemiluminescence (Thermo Scientific). For chondroitin-sulfate proteoglycan detection, lysates were treated according to the detailed deglycosylation protocol from Mariano Viapiano Ph.D. lab website (Massey et al., 2008). Briefly, 50-100μg of protein lysate was treated for 6-8hrs with chABC (100 μU/mL; R&D systems) before being subjected to traditional western blotting, probed with anti-chondroitin-4-sulfate (1:1000; Millipore: MAB2030) or anti-chondroitin-6-sulfate (1:1000; Millipore: MAB2035). Protein expression was quantified with ImageJ densitometry and normalized to total protein as measured by ponceau staining.

### Sympathetic outgrowth assay

Cultures of dissociated sympathetic neurons were prepared from superior cervical ganglia (SCG) of newborn rats as described (Dziennis & Habecker, 2003). Cells were pre-plated for 1 hour to remove non-neuronal cells, and then 5,000 neurons/well were plated onto a 96 well plate (TPP) coated with (poly-L-lysine PLL, 0.01%, Sigma-Aldrich) and either laminin (10 μg/mL, Trevigen), laminin and CSPGs (2μg/mL; Millipore), or laminin and CSPGs pre-treated with ARSB (0.3 or 0.6μg/mL; R&D systems) for 6-8hrs. Neurons were cultured in serum free C2 medium (Pellegrino, Parrish, Zigmond, & Habecker, 2011) supplemented with 10 ng/mL NGF (Alomone Labs), 100 U/mL penicillin G, and 100 μg/mL streptomycin sulfate (Invitrogen). Live cell imaging was carried out using an Incucyte Zoom microscope (Essen BioScience), with 20x phase images acquired every 2hrs over a 40hr period. Neurite length was measured using Cell Player Neurotrack software (Essen BioScience) and was used to calculate the neurite growth rate.

### Myocardial ischemia/reperfusion surgery

Anesthesia was induced with 4% isoflurane and maintained with 2% isoflurane. Mice were restrained supine, intubated, and mechanically ventilated. Core body temperature was monitored by a rectal probe and maintained at 37°C throughout the surgery. The left anterior descending coronary artery (LAD) was reversibly ligated for 40 minutes and then reperfused by release of the ligature. LAD occlusion was verified by a persistent S-T wave elevation, region specific cyanosis, and wall motion abnormalities. Reperfusion was confirmed by return of S-T wave to baseline level and re-coloration of ventricle region distal to occlusion(Gardner & Habecker, 2013; Parrish et al., 2010).

### Explant Co-culture Assay

Explants were generated as previously described (Gardner & Habecker, 2013). Briefly, newborn mouse superior cervical ganglia were explanted and placed into culture 1mm from left ventricle tissue – either cardiac scar tissue (10-14 days post-MI) or unoperated control tissue. Cardiac tissue from a single animal was split in half and cultured with the left or right ganglia from a single animal, this enabled tissue from the same mice to be treated with either vehicle (5% DMSO) or ARSB to remove 4S-CS GAGs. The ganglia and the cardiac tissue were co-cultured inside a Matrigel bubble surrounded by C2 media supplemented with 2ng/mL NGF. ARSB was added directly to the media (0.6 μg/mL) at the time of plating and again 24hrs later. At 48hr the explants were imaged by phase microscopy with a Keyence BZ-X microscope. Neurite length from the edge of the ganglia to the most distal tip of visible neurites was measured using ImageJ.

### siRNA pool knockdown efficiency screen

A pool of 4 siRNAs targeting the *Chst15* gene were purchased from Horizon Discovery (formerly GE Dharmacon) and were tested for their efficacy in reducing gene expression in C2C12 myoblasts. C2C12 myoblasts were plated on 12-well plates coated with collagen and were transfected with the various targeting and control siRNAs using the Dharmafect transfection reagent (3μL of Dharmafect reagent per well with 120nM siRNA). The next day wells were split using Versene (Gibco), 1/3 of the cells were collected for a 24-hour time point and the other 2/3 were split into 2-wells for 48hr and 72hr knockdown timepoints. Cells were processed using an RNA Mini-kit (Qiagen) to purify RNA. 1μg of RNA was loaded into the cDNA reaction with the iScript cDNA synthesis kit. Gene expression was examined with multiplexed Taqman probes targeting *Chst15* and *Gapdh* using 2μL of cDNA template and Taqman reagents and measured with an ABI7500 Thermocycler. The delta delta ct method was used to calculate knockdown efficiency. Once an effective siRNA against *Chst15* was identified, knockdown efficiency was measured at the protein level with C2C12 myoblasts 48hr post-knockdown. CHST15 expression was assessed via western blot with a CHST15 antibody (1:1000; Proteintech: 14298-1-AP) normalized to GAPDH (1:1000; Thermo Scientific: MA1-16757); siRNA *Chst15-2* was selected for larger scale production and used for in vivo studies.

### In vivo siRNA treatment

After MI surgery mice were treated with 100μg of siRNA targeting *Chst15* or a non-targeting control. siRNA was delivered systemically via tail-vein injection on days 3, 5 and 7 following MI. Tissue was collected at day 10 post-MI for western blot analysis of CSPG sulfation and the sympathetic neuron marker TH. siRNA used for this experiment was custom synthesized by Horizon Discovery; siAccell *in vivo* formulation.

### NE content by High-Performance Liquid Chromatography

NE levels in heart tissue were measured using HPLC with electrochemical detection (Li, Knowlton, Van Winkle, & Habecker, 2004). Frozen, pulverized tissue was weighed, and then homogenized in 0.2M perchloric acid and 0.5 mM dihydroxybenzylamine (DHBA). The tissue was refrigerated for 1hr and centrifuged at 14,000 rpm for 4min. Supernatant was adsorbed on alumina, followed by 15min of tumbling. The alumina was washed twice with 1.0mL of H2O with centrifugations in between. The NE was desorbed from the alumina with 150 ml of 0.1M perchloric acid. 50μL aliquots were fractionated by reversed-phase HPLC (C18; 5 mm particle size, Rainin) using a mobile phase containing 75mM sodium phosphate (pH 3.0), 360mg l 1 of sodium octane sulfonate, 100mL l 1 triethylamine and 3.0% acetonitrile. A coulometric detector (Coulchem, ESA) was used to detect and quantify NE and DHBA. NE standards (0.5 mM) were processed in parallel with tissue samples and interspersed throughout the HPLC run. Retention time for NE was 5.0min and for DHBA was 8.5min.

### Immunohistochemistry

Tyrosine hydroxylase (TH; sympathetic nerve fibers) and fibrinogen (Fib; infarct/scar) staining was carried out as described previously (Gardner et al., 2015). Tissue was collected 10 days after surgery, fixed in 4% paraformaldehyde, frozen and 12μm sections generated. To reduce autofluorescence sections were rinse 3 x 10 minutes in 10mg/mL Sodium Borohydride and rinsed for 3 x 10 minutes in PBS. Slides were placed in 2% BSA, 0.3% Triton X-100 in PBS for 1 hour and then incubated with rabbit anti-TH (1:1000; Millipore: AB1542) and sheep anti-fibrinogen (1:300; BioRad: 4400-8004) overnight. The following day the slides were incubated with Alexa-Fluor IgG-specific antibodies (Molecular Probes, 1:1000) for 1.5 hr and rinsed 3 x 10 minutes in PBS. Background autofluorescence was reduced further with a 30min incubation in 10mM CuSO4 (diluted in 50mM Ammonium Acetate). Following this, slides were rinsed 3 x 10 minutes in PBS before mounting in 1:1 glycerol:PBS and visualized by fluorescence microscopy. Threshold image analysis of TH staining has been described previously (Gardner et al., 2015) but briefly, the threshold function in ImageJ was used to generate black and white images discriminating TH+ nerves for 6 sections spanning 200μm of the infarct/scar or peri-infarct region from each heart. Percent area TH+ fiber density (20x field of view) was quantified within the infarct and the area immediately adjacent to the infarct (peri-infarct). Images were acquired with a Keyence BZ-X 800 microscope.

### Quantification of Cardiac Scar (infarct) size

Infarct size was determined 10 days after myocardial ischemia-reperfusion injury and tissue was prepared for immunohistochemistry as previous mentioned omitting the steps to reduce autofluorescence. Autofluorescence in the GFP channel was used to image the infarct size. Scar tissue is notably lacking autofluorescence enabling easy identification of the infarct. Images were acquired with a Keyence BZ-X 800 microscope at 2x magnification and analyzed using Image J freehand selection tool. Left ventricle (LV) and infarct was outlined, measured and the percent area of cardiac scar was determined by (infarct area/ LV area) x 100. The scar was imaged in 6 sections across 200μm of the infarct as previously described(Gardner et al., 2015).

### Arrhythmia Assessment

Anesthesia was induced with 4% isoflurane and maintained with 2% isoflurane in day 10 post-MI animals treated with siRNA. ECG leads were connected to monitor arrhythmias and animals were maintained at 37°C throughout the analysis. All parameters were monitored with Powerlab LabChart software (AD Instruments). A 30-minute baseline was used to assess spontaneous arrhythmia susceptibility prior to administration of *ββ*-agonist Isoproterenol (50μg) and caffeine (3mg) as described previously(L. Wang et al., 2014). Arrhythmias were measured for 30 minutes following drug administration and scored according to the modified Lambeth conventions (Curtis et al., 2013) on a scale of 0-4. Individual animals received a single score based on the most severe arrhythmia observed. 0 indicates no arrhythmia. 1 indicates 1-2 premature ventricular contractions (PVCs) followed by normal sinus rhythm of at least 2 beats. 2 indicates bigeminy (1 PVC followed by one normal sinus beat, repeating for 4 or more continuous cycles) or salvo (3-5 PVCs in a row). 3 indicates non-sustained ventricular tachycardia (nsVT) defined as 6 or more PVCs in a row lasting less than 30 seconds. 4 indicates sustained VT (>30 seconds) or Torsades de Pointes.

### Statistics

Student’s t-test was used for comparisons of just two samples. Data with more than two groups were analyzed by one-way ANOVA using the Tukey post-hoc test to compare all conditions or the Dunnet’s post-test when comparing to a single control group. Data with multiple variables was analyzed by two-way ANOVA. All statistical analyses were carried out using Prism 9.

## Acknowledgements

The authors would like to thank Kevin Wright Ph.D for crucial comments on the manuscript and Ryan Gardner Ph.D for critical insight during the early stages and development of this project. We would also like to thank Tammi Howard for early technical assistance on this project. This work was supported by NIH F31HL152490 (MRB), AHA 20PRE35210768 (MRB), the Steinberg Endowment for Graduate Education (MRB), Portland State University EXITO Scholars Program funds (SS), and NIH R01 HL093056 (BAH).

## Supplemental Data

**Figure S1:**
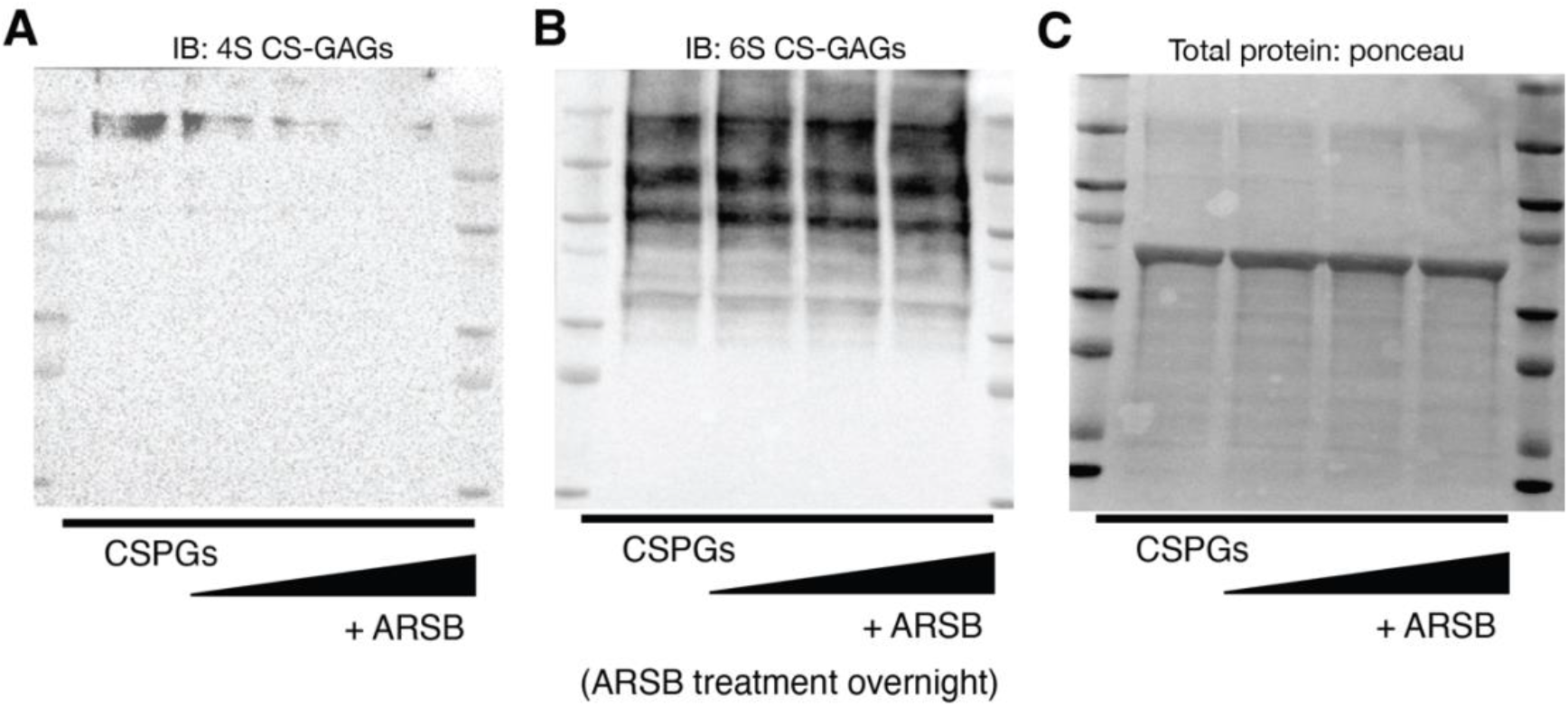
ARSB removes 4S CS-GAGs and leaves 6S CS-GAGs intact. **(A)** Western blot of 4S content using purified CSPGs upon treatment of increasing concentrations of ARSB; vehicle, 0.3μg/mL, 0.6μg/mL, and 1.2μg/mL respectively left to right. **(B)** Western blot of 6S using purified CSPGs upon treatment of increasing concentrations of ARSB; vehicle, 0.3μg/mL, 0.6μg/mL, and 1.2μg/mL respectively left to right. **(C)** Total protein loaded of purified CSPGs treated with increasing concentrations of ARSB; vehicle, 0.3μg/mL, 0.6μg/mL, and 1.2μg/mL respectively left to right.

**Figure S2:**
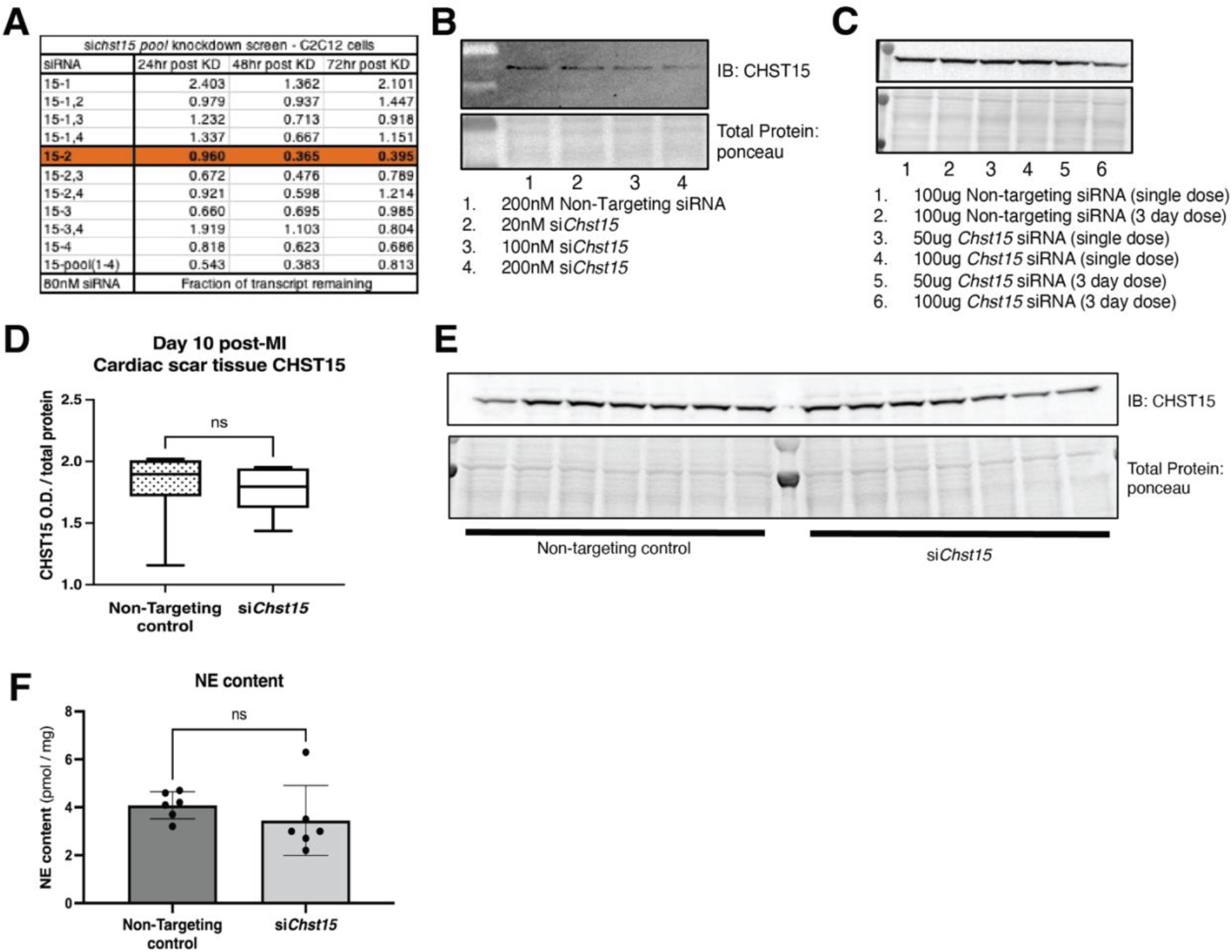
Identification of an effective siRNA against *Chst15*. **(A)** qPCR knockdown of siRNA pool in C2C12 cells in 3 days after knockdown. Data shown fraction of transcript remaining, si*Chst15-2* was most effective in knockdown of *Chst15*. This transcript was selected for *in-vivo* studies **(B)** Western blot of CHST15 protein knockdown in C2C12 cells to confirm efficacy of si*Chst15-2*, comparison to Non-Targeting controls **(C)** Tail vein injection in mouse to determine ideal dosing for *in-vivo* si*Chst15* knockdown. Western blot of tail vein injection dosing trial for si*Chst15* with either 1 day or 3 days of injections. CHST15 protein in left ventricle (LV) 48hr after final tail vein injection, comparison to Non-Targeting controls, all unoperated (non-MI) animals. **(D)** Full siRNA CHST15 experimental trial, CHST15 protein expression on D10 post MI quantified, siRNA injection D3, 5, 7 post-MI, n=7 animals per treatment group, statistics; student t-test (Welch’s test), ns – not significant. **(E)** Western blot of CHST15 protein expression D10 post-MI in siRNA treated animals. **(F)** NE content in the cardiac scar following siRNA treatment. Quantification of n=6 animals for non-targeting controls and *Chst15 siRNA* treatment. Statistics; student t-test (Welch’s test), n.s.-not significant.

